# Benchmarking of automated cancer cell annotation methods for scRNA-seq data reveals Consensus annotation as the preferred method

**DOI:** 10.1101/2025.01.13.632750

**Authors:** Tamim Abdelaal, Christiaan Klijn, Matt Hancock

**Affiliations:** Discovery Data Science, Genmab B.V., Utrecht, The Netherlands

## Abstract

Targeted cancer therapies have shown therapeutic advantages due to tumor-specific drug activity. Single-cell RNA-sequencing has been widely used in cancer studies to define different cellular identities. However, accurate identification of tumor vs other normal cells is essential to define novel tumor-specific targets. Recent methods have been developed to perform the task of tumor cell annotation, which can be divided into two categories: CNV-based methods, which use transcriptome measurements to infer copy number variations and identify cells with alterations as tumor, and Reference-based methods which train a classifier using previously annotated tumor data and use it to annotate new datasets. We benchmarked the state-of-the-art method of each category, SCEVAN and scATOMIC, respectively, together with Consensus annotation method where a cell is considered tumor if both methods agree on that. Across 20 cancer datasets spanning 9 cancer types with a total of 379 samples, the Consensus annotation outperformed other methods in terms of precision score. SCEVAN works well when clear CNVs are detected, otherwise cells are randomly split between normal and tumor producing many false positives. While scATOMIC efficiently detects all normal cells except normal epithelial cells, where all epithelial cells (normal and malignant) are considered cancerous. The Consensus annotation method overcomes the limitations of both methods, being able to detect normal epithelial cells together with other normal cell types as non-tumor, also annotating malignant epithelial cells as tumor even with weak CNVs. This produces overall higher precision with the least number of false positives, leading to confident tumor-specific potential therapeutic targets. Implementation is available in the MACE R package through GitLab (https://gitlab.com/genmab-public/mace/).

## 1 Introduction

Understanding the complex interactions within tumor microenvironments (TMEs) is crucial for elucidating the mechanisms of cancer proliferation, progression, and therapeutic resistance [1-3]. Single-cell RNA sequencing (scRNA-seq) has emerged as a powerful tool to dissect these complicated systems by providing comprehensive transcriptomic profiles of individual cells within a tumor, facilitating the detailed study of cellular heterogeneity and the dynamic interplay between malignant cells and their surrounding immune and stromal components [4-6].

Accurate cell type annotation in scRNA-seq data is pivotal for deriving meaningful biological insights, which can be performed manually or automated [7]. Traditional manual annotation methods, which rely heavily on pre-existing knowledge and canonical gene markers, have significant limitations, including subjectivity and incomplete markers knowledge. On the other hand, automated classifier-based methods have been developed, offering more consistent and scalable annotations [8, 9]. In the context of cancer, it is important to dissect malignant tumor cells from the surrounding normal immune and stromal cells [10]. While designing targeted cancer therapies, accurate identification of malignant tumor cells is essential to enable tumor-specific drug activity, avoiding damaging or depletion of other normal cells in the TME.

Malignancy refers to the presence of cancerous cells that exhibit uncontrolled growth, invasion into surrounding tissues, and the potential to metastasize to distant organs. Defining malignant cells in single-cell genomics poses significant challenges due to the heterogeneity within tumors and the dynamic nature of cancer progression. Malignant cells often exhibit genomic instability, characterized by mutations, copy number variations (CNVs), and other genetic alterations that differentiate them from normal cells [11]. However, the tumor microenvironment (TME) is composed of a complex mixture of malignant cells, immune cells, stromal cells, and other non-cancerous components, making it difficult to distinguish malignant cells with high precision [12]. Additionally, the variability in gene expression profiles among malignant cells, driven by factors such as clonal evolution and environmental influences, further complicates their identification [13].

Several computational methods have been developed to identify tumor cells using scRNA-seq data, which can be divided into two broad categories. First, methods relying on training a classifier using previously annotated cancer reference datasets, hence Reference-based methods. Next, using the trained classifier model to annotate new datasets. Reference-based methods include Seurat [14], scmap [15], SingleR [6], CHETAH [16], scType [17] and SingleCellNet [18] among others. Although these methods generally perform the task of cell identification well, those are not designed to differentiate the complex TME of cancer studies. Therefore, a recent method scATOMIC [19] was designed specifically to identify cells from cancer scRNA-seq datasets, showing overall superior performance especially for the cancer cells compartment. Thus, scATOMIC represents the current state-of-the-art Reference-based method.

The second category of methods identifying tumor cells are relying on inference of copy number variation (CNV) from scRNA-seq gene expression data, hence CNV-based methods. Various methods including inferCNV [5], CopyKAT [20] and SCEVAN [21], all depending on the assumption of defining tumor cells having genomic instabilities compared to a normal diploid genome for normal cells. Recently, SCEVAN showed superior performance to other methods, representing the current state-of-the-art CNV-based method. Also, SCEVAN can independently detect a set of confident normal cells to be used to define the normal diploid genomic profile, i.e. it does not require a set of normal cells as an extra input.

Currently, it is not clear which category of methods is more effective when identifying malignant tumor cells in scRNA-seq data analysis. Further, an independent evaluation is required to compare methods across various cancer types, guiding users to which method fits best their data, and guiding further developments in the field.

In this study, we benchmarked the state-of-the-art method from each category: Reference-based (scATOMIC) and CNV-based (SCEVAN), for the task of precise identification of tumor vs normal cells using scRNA-seq data. The evaluation was performed using 20 cancer scRNA-seq studies covering 9 different cancer types spanning a total of 379 patient samples. As results show, the performance of the two methods varied across cancer types, hence, we designed a Consensus annotation method which carried the benefits and overcomes the limitations of both Reference-based and CNV-based methods. The Consensus annotation method showed superior tumor cell annotation performance producing overall higher precision across all datasets.

## 2 Methods

### 2.1 Tumor cell identification methods

We evaluated the state-of-the-art methods scATOMIC [19], denoted as the Reference-based method, and SCEVAN [21] denoted as the CNV-based method, for **Ma**lignant **Ce**ll (MACE) annotation. Where the MACE annotation is defined as the task of annotating tumor vs normal cells using scRNA-seq dataset. Both methods are publicly available as R packages, and were applied using their default settings, unless otherwise stated. During the benchmarking, we provided the input scRNA-seq datasets as raw count data. The annotation was performed one patient sample at a time to avoid differences across individuals. Finally, the predicted annotation by each method is evaluated against the ground-truth cell type annotation provided by the authors of each dataset/study (Fig. 1A).

**Fig. 1.**
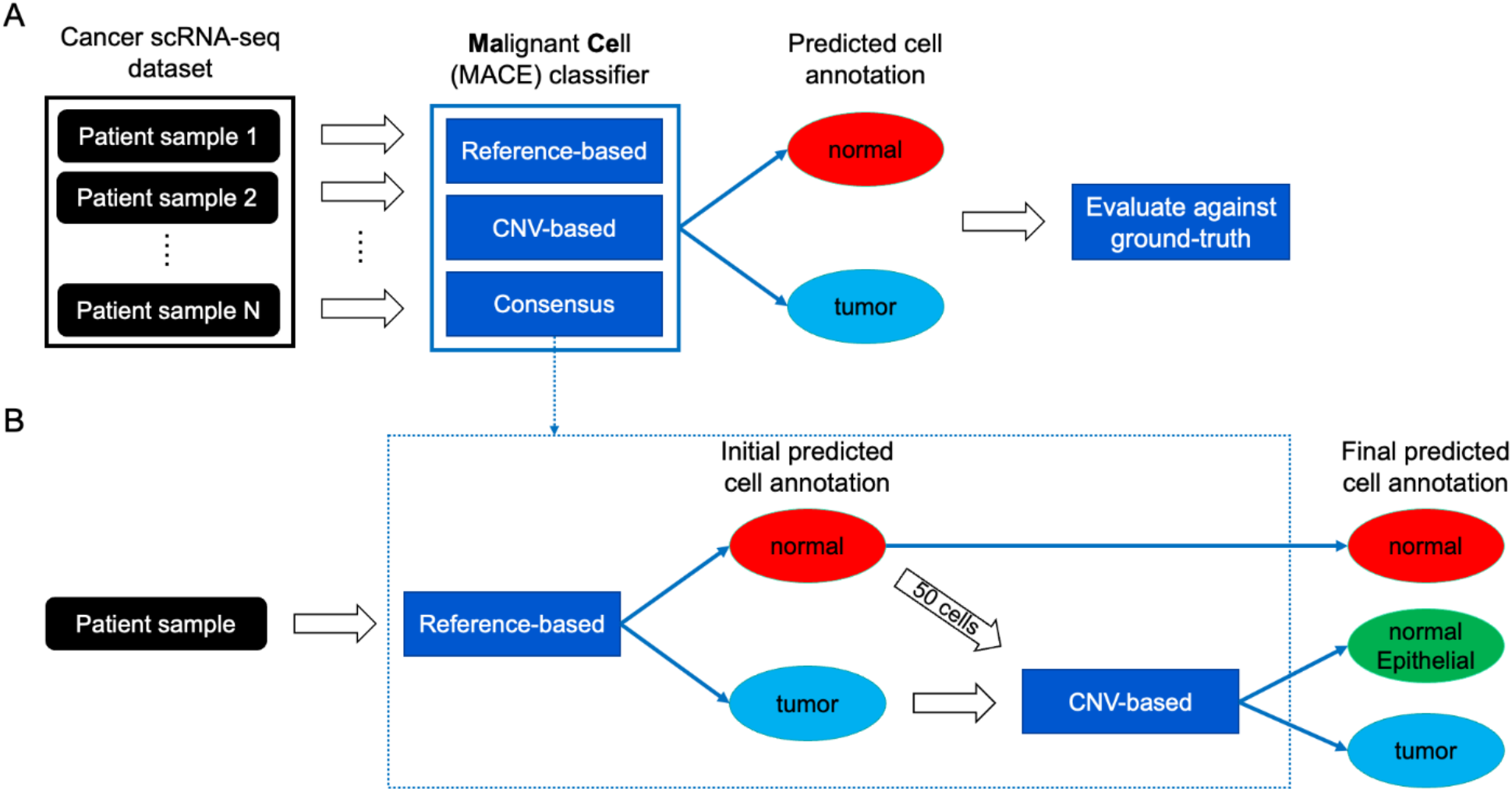
(A) Benchmarking pipeline. Cancer scRNA-seq dataset is provided as input raw count data to each **Ma**lignant **Ce**ll (MACE) classification method per patient samples. Each method predicts tumor vs normal cell annotation which is further evaluated against the ground-truth cell type annotation of each dataset. **(B) Consensus annotation pipeline**. Combination of the Reference-based and the CNV-based methods, the Reference-based method generates the initial cell annotation, while the CNV-based refine the initial tumor annotation further into tumor and normal epithelial cells.

The Reference-based method, scATOMIC, provides a hierarchy of cell type annotations. For the task of predicting tumor vs normal cells, we stop at a high level in the hierarchy at layer 2. Cells classified as ‘Non Stromal Cell’ are considered ‘tumor’ cells, other cell type annotations are considered ‘normal’ cells, and any unclassified cell annotation is left as ‘unclassified’. The CNV-based method, SCEVAN, directly provides a class annotation for each cell either being ‘tumor’ or ‘normal’. Both methods filter out low-quality cells during their preprocessing, these left out cells are annotated as ‘filtered’. These filtered cells were also left out during the evaluation against the ground-truth annotation.

We added a third method combining the Reference-based and the CNV-based methods, denoted as the Consensus annotation method (Fig. 1B). As suggested by the method name, the Consensus annotation identify a tumor cell where both Reference-based and CNV-based methods agree on that ‘tumor’ annotation. The detailed steps of the Consensus annotation are: 1) Performing the Reference-based annotation on all cells within the input sample, to obtain an initial tumor vs normal cell annotation, where the tumor annotation represents the epithelial cells. 2) Performing the CNV-based annotation only on the tumor cells identified by the Reference-based method to further split them into normal epithelial cells, having normal diploid genomic profiles, and tumor cells with abnormal genomic CNV profiles. Here, we provide the CNV-based method with a sample of 50 cells from the ‘normal’ annotation by the Reference-based method, representing the reference normal diploid genomic profile. This is mainly to speed-up the computation time and skip the internal step of the CNV-based method to identify these normal cells by itself. 3) The final annotations by the Consensus method are: ‘normal’ cells identified by the Reference-based method, ‘normal Epithelial’ cells identified as tumor cells by the Reference-based method and redefined as normal cells by the CNV-based method, and ‘tumor’ cells identified as tumor cells by both methods. Filtered out low-quality cells are annotated as ‘filtered’.

### 2.2 Datasets

We used a total of 20 cancer scRNA-seq datasets to evaluate all methods. Datasets are covering 9 different cancer types having a combined number of 379 cancer samples (summarized in **Table** 1). All datasets have been obtained from the original publications, including the raw count gene expression data, along with the cell type annotation provided by the authors of each study. Only cancer samples were retained from each study, blood or adjacent normal samples were excluded.

**Table 1.** Overview of the datasets used during this study.

| **Dataset** | **No. of samples** | **No. of cells** | **Cancer type** | **Reference** |
| --- | --- | --- | --- | --- |
| Elyada_et_al_PDAC | 6 | 16,836 | Pancreas | [22] |
| Hwang_et_al_PDAC | 18 | 108,964 |  | [23] |
| Peng_et_al_PDAC | 24 | 41,986 |  | [24] |
| Steele_et_al_PDAC | 17 | 48,453 |  | [25] |
| Horny_et_al_HNSC | 21 | 40,102 | Head and Neck | [26] |
| Kurten_et_al_HNSC | 33 | 113,579 |  | [27] |
| Puram_et_al_HNSC | 18 | 5,740 |  | [28] |
| Gao_et_al_Breast | 8 | 16,633 | Breast | [20] |
| Qian_et_al_Breast | 14 | 44,024 |  | [29] |
| Wu_et_al_Breast | 20 | 81,646 |  | [30] |
| Pelka_et_al_CRC | 60 | 236,679 | Colorectal | [31] |
| Qian_et_al_CRC | 7 | 44,684 |  | [29] |
| Lee_et_al_CRC | 23 | 47,285 |  | [32] |
| Khaliq_et_al_CRC | 16 | 49,859 |  | [33] |
| Kim_et_al_Lung | 32 | 107,761 | Lung | [34] |
| Qian_et_al_Lung | 8 | 93,575 |  | [29] |
| Jerby-Arnon_et_al_Skin | 23 | 5,735 | Skin | [35] |
| Ma_et_al_Liver | 15 | 8,855 | Liver | [36] |
| Bi_et_al_Kidney | 8 | 34,326 | Kidney | [37] |
| Chen_et_al_Bladder | 8 | 37,039 | Bladder | [38] |

### 2.3 Data preprocessing

The raw count gene expression data was used as input to all methods. For the Reference-based method, scATOMIC, cells expressing less than 200 genes or having more than 25% mitochondrial genes were filtered out. For the CNV-based method, SCEVAN, cells expressing less than 200 genes were filtered out, as well as genes expressed in less than 10% of the cells.

### 2.4 Evaluation

We considered the tumor vs normal annotation task as a binary classification, with ‘tumor’ being our positive label. Therefore, we evaluated the prediction performance of each method using the Precision score:

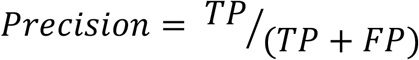

Where TP is the true positives representing actual tumor cells correctly annotated as tumor cells, and FP is false positives representing normal cells wrongly annotated as tumor cells. Since the annotation was performed per patient sample, the precision score was calculated per patient sample as well.

### 2.5 Implementation details

For the Reference-based method, scATOMIC, we applied the *run_scATOMIC* and *create_summary_matrix* functions, while setting the input parameters *fine_grained_T, use_CNVs*, and *modify_results* to *False*. We obtained the cell annotations from the ‘layer_2’ output. For the CNV-based method, SCEVAN, we applied the *pipelineCNA* function, while setting the input parameters *SUBCLONES* and *plotTree* to *False*. We obtained the cell annotations from the ‘class’ output.

We developed wrapper functions, compatible with Seurat objects, to perform the Reference-based, CNV-based and Consensus tumor vs normal annotation. The implementation is available as the MACE R package through GitLab (https://gitlab.com/genmab-public/mace/).

## 3 Results

### 3.1 Annotation performance varies per cancer type

We started by benchmarking the Reference-based and the CNV-based methods for the task of annotating tumor vs normal cells, using all 20 scRNA-seq datasets covering 9 different cancer types. We compared the predicted cell annotations provided by each method against the cell type annotation provided by the authors of each study and calculated the precision score per patient sample. From the authors annotation, we considered the malignant/cancer cells population as the ground-truth ‘tumor’ and all other cell populations as ‘normal’. We based the benchmarking results on the precision score, although other classification metrics could be used. However, in the context of targeted therapy on cancer cells, it is more important to define a purified set of tumor cells, instead of annotating more tumor cells with contamination from other normal cells. Therefore, we aimed to find the method producing the best precision score to control and minimize the false positive predictions.

Based on the precision score, we observed that results vary across cancer types. The Reference-based method produced better average performance for Head and Neck, Breast, Colorectal, Lung and Kidney cancers. Where more samples had higher precision score for the Reference-based method compared to the CNV-based method (Fig. 2). However, for Pancreas and Liver cancers, the CNV-based method produced higher performance. Finally, both methods had comparable performance for Skin and Bladder cancers. Moreover, the precision score produced in most samples is relatively high, however, large performance variability is observed for the CNV-based method for Head and Neck, Breast, Lung and Kidney cancers. Similarly, for the Reference-based method when applied on Pancreatic cancer samples. Overall, this shows that on average both methods can produce good annotation performance depending on the cancer type.

**Fig. 2.**
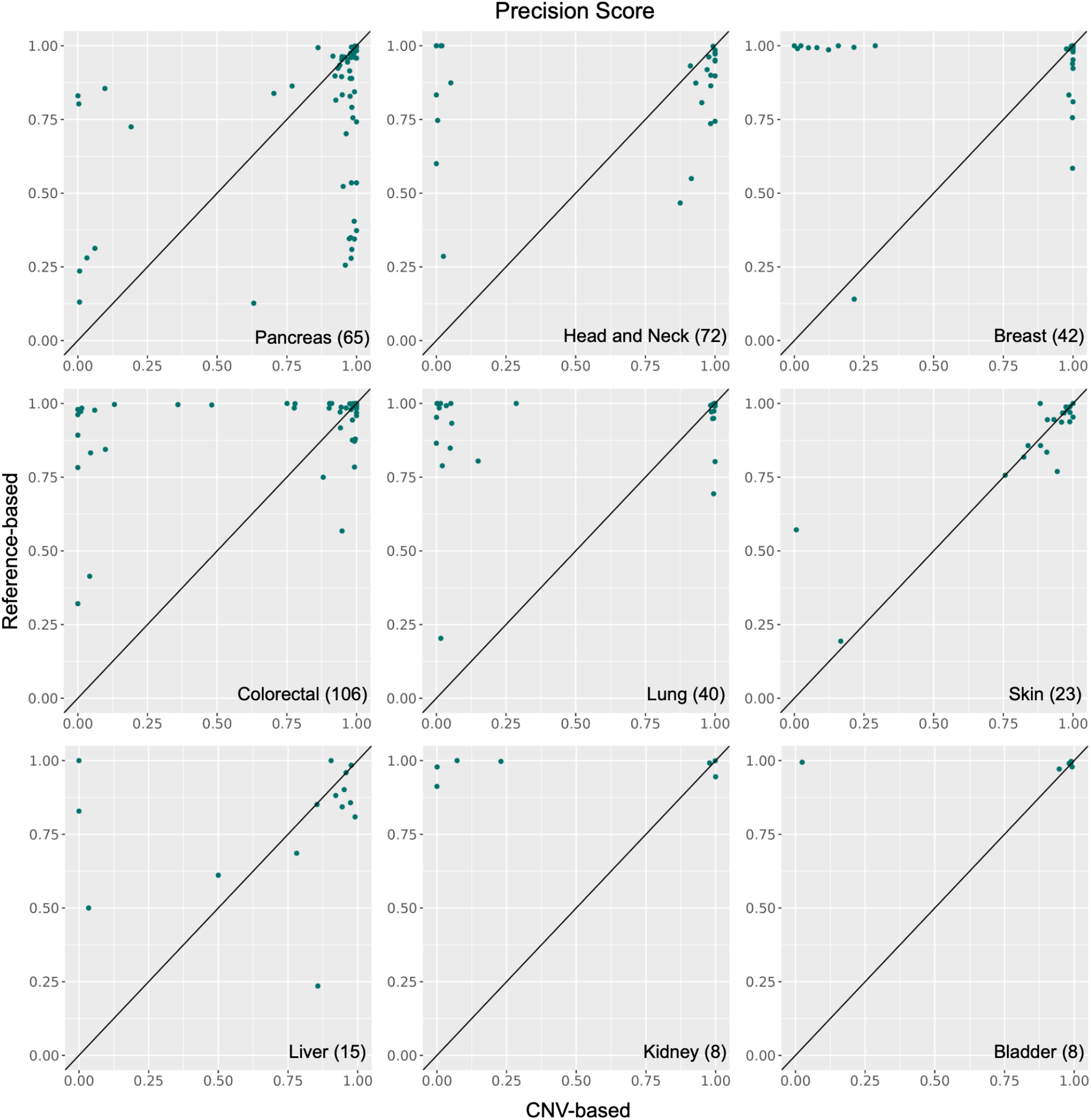
Benchmarking results for the Reference-based vs the CNV-based methods. Scatterplots showing the precision score of the Reference-based and the CNV-based methods across 9 cancer types. Each point represents a patient sample, where more points on the upper triangle shows higher Reference-based performance, while more points in the lower triangle shows higher CNV-based performance.

### 3.2 Consensus annotation combines the power of other methods

We investigated the limitations of the Reference-based and the CNV-based methods, by examining when and why cells are wrongly annotated. Taking the Peng_et_al_PDAC dataset as an example, where the Reference-based method performed poorly. According to the authors cell type annotation, two ductal cell populations were identified, where the Ductal cell type 2 is considered the malignant population based on copy number variation profile (Fig. 3A). Comparing with the Reference-based tumor annotation, all immune and stromal cell types were correctly annotated as ‘normal’ cells. However, together with the Ductal cell type 2, the Reference-based method included the Acinar cells and Ductal cell type 1 as ‘tumor’ cells, which are considered mis-annotations (Fig. 3A). The latter two populations were correctly separated from the ‘tumor’ cells by the CNV-based method, which more precisely annotated only the Ductal cell type 2 as ‘tumor’ (Fig. 3A).

**Fig. 3.**
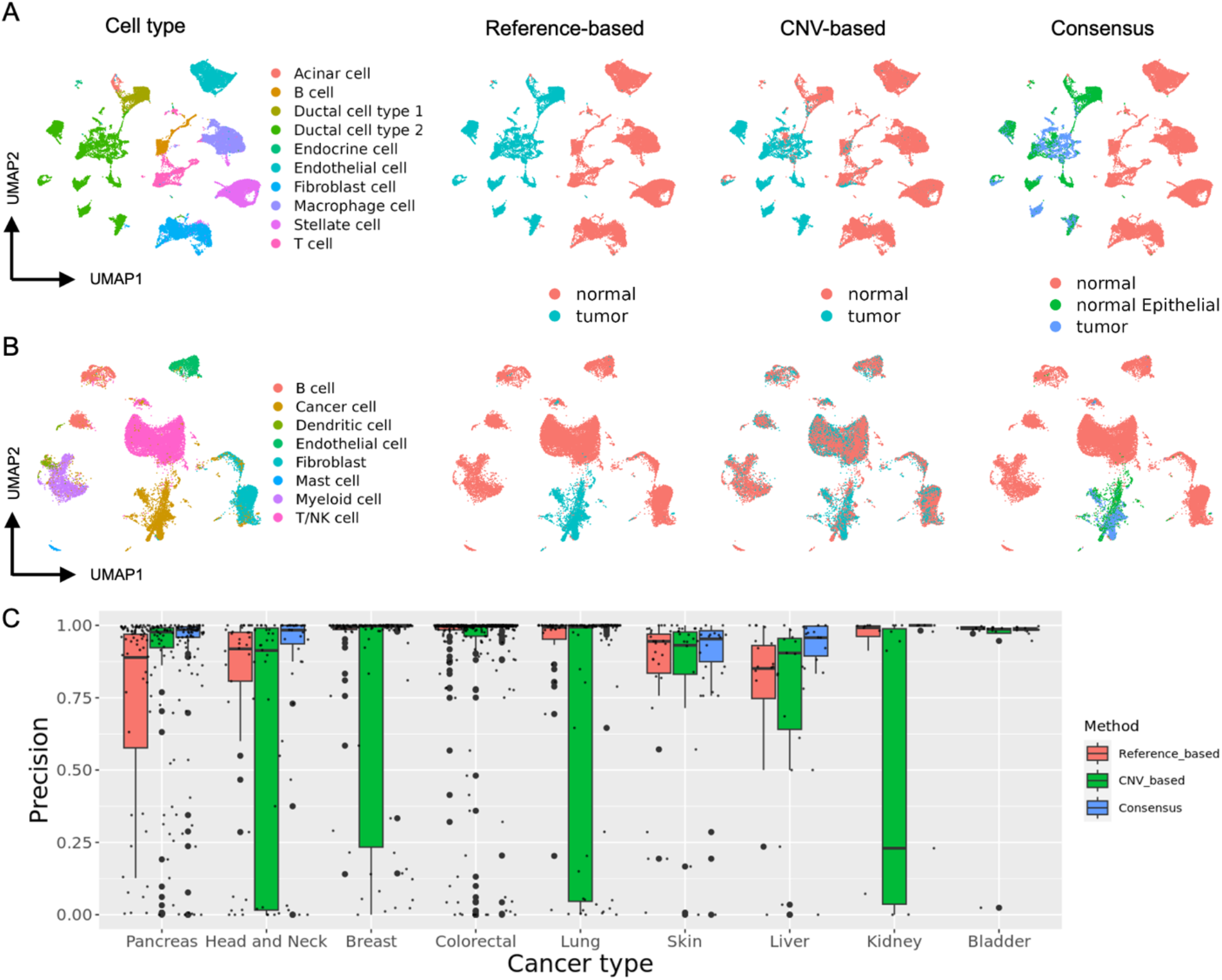
**(A-B)** UMAP visualization [39] of **(A)** the Peng_et_al_PDAC and **(B)** the Qian_et_al_Breast datasets colored according to the authors cell type annotation, in comparison with the MACE annotation produced by the Reference-based, the CNV-based and the Consensus methods. **(C)** Boxplot comparing the performance of all methods, based on the precision score, showing that the Consensus method produced the most precise annotation across 9 different cancer types.

On the other hand, investigating the Qian_et_al_Breast dataset as an example of poor performance for the CNV-based method. The authors annotated the tumor cells as the Cancer cell population (Fig. 3B). The CNV-based method almost randomly splits all cells between tumor and normal cells, including many wrong ‘tumor’ annotations from immune and stromal cell types (Fig. 3B). However, the Reference-based method performed well by segregating the Cancer cell population as ‘tumor’ from all other ‘normal’ cells (Fig. 3B).

Building on this information, we combined both methods into Consensus annotation to retain the power and overcome the limitation of each method. The motivation is to have a precise tumor vs normal annotation method, which: 1) can accurately separate potential epithelial cancer cells from all other normal cells in the TME, and 2) can further separate the epithelial cells compartment into malignant/tumor and normal cells based on CNV profiles, thus retaining the normal epithelial cells as non-malignant. The Consensus method uses the Reference-based method to achieve the first task of identifying epithelial vs other normal cells. Followed by the CNV-based method applied only on those epithelial cells (see Methods), together with examples of normal cells for normal CNV diploidy profile, to further separate normal from malignant epithelial cells. For the Peng_et_al_PDAC dataset, the Consensus method was able to separate the Acinar cells and the Ductal cell type 1 as ‘normal’, benefiting from the CNV-based method (Fig. 3A). Further, for the Qian_et_al_Breast dataset, the Consensus method annotated tumor cells only from the Cancer cell population, benefiting from the Reference-based method. Also, it reverts all the wrong ‘tumor’ annotations made by the CNV-based method to ‘normal’ cells (Fig. 3B). Across all datasets, the Consensus annotation method outperforms other (separately applied) methods, producing the highest average precision score across all cancer types (Fig. 3C). This shows that the Consensus method combines the benefits of both Reference-based and CNV-based methods, offering a new solution to precisely annotate tumor cells using scRNA-seq data.

### 3.3 Consensus method defines tumor cells with valid CNV profiles

Next to producing the best annotation performance, we aimed to validate that the final tumor epithelial cells identified by the Consensus method have valid CNVs, in comparison to their normal epithelial counterpart. Inspecting the CNV profiles of the Consensus method indeed showed clear CNV profile differences between tumor and normal epithelial cells. As observed by example patient samples from the Peng_et_al_PDAC and Qian_et_al_Breast datasets, tumor epithelial cells are experiencing various genomic deletions (loss) and duplications (gain) across multiple genomic locations (Fig. 4, left column). In contrast, the normal epithelial cells show a homogenous diploid CNV profile as expected.

**Fig. 4.**
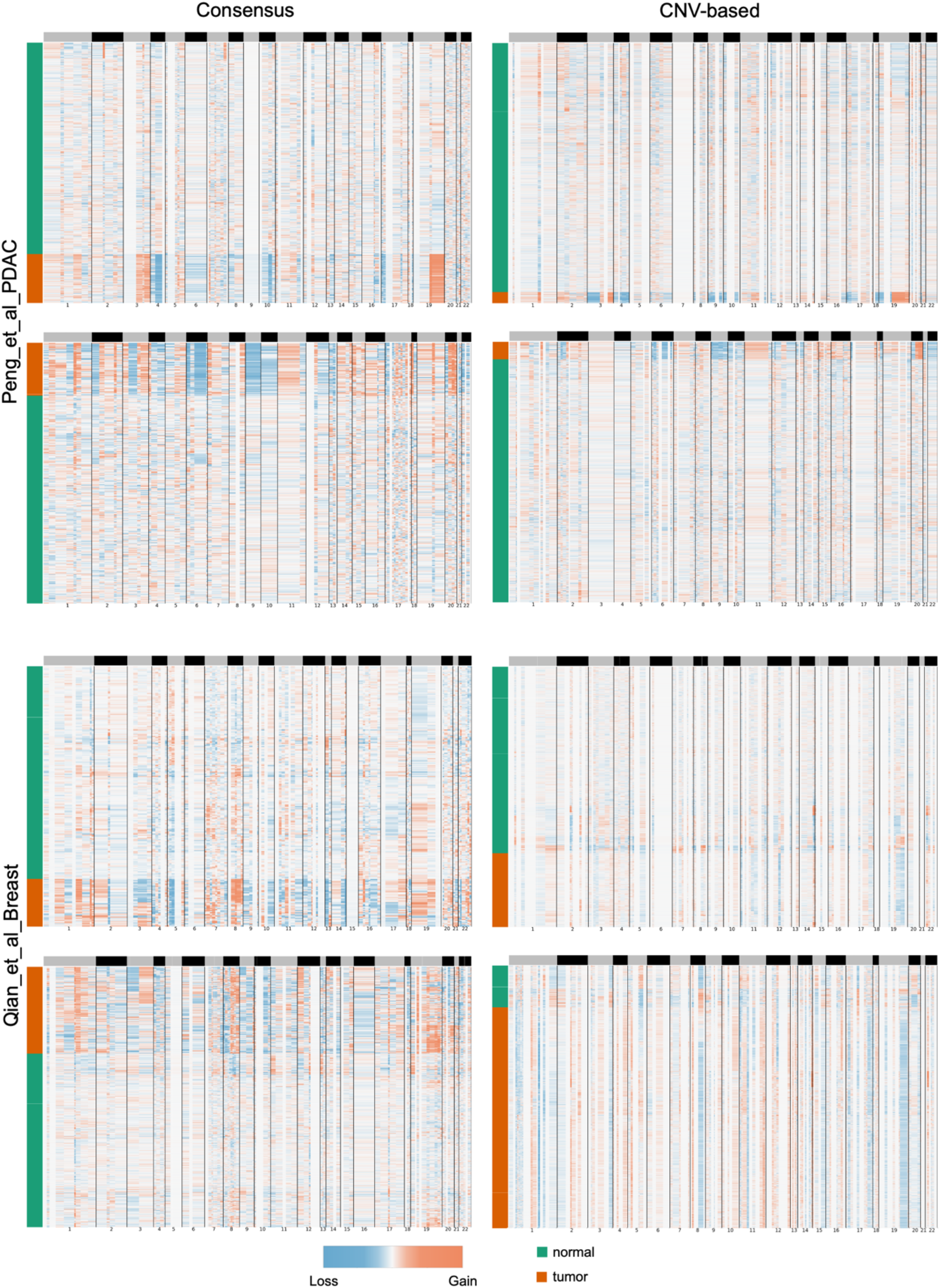
CNV profiles of tumor vs normal cells. Heatmaps showing the CNV profiles produced by the Consensus method (left column) and the CNV-based method (right column) across samples from the Peng_et_al_PDAC dataset (top two rows) and the Qian_et_al_Breast dataset (bottom two rows). Each heatmap shows cells x genomic locations (rows x columns), colored according to the copy number variation (blue=loss, red=gain, white=normal diploid). The vertical sidebar to each heatmap indicates the tumor vs normal predicted annotation.

More importantly, we compared the CNV profiles produced by the Consensus method with the CNV profiles produced by the CNV-based method for the same samples (Fig. 4, right column). Recall that the CNV-based method is applied across all cells, not only the epithelial cells compartment. For the Peng_et_al_PDAC samples, the tumor cells annotated by the CNV-based method had clear CNV aberrations compared to normal cells. These clear CNVs suggest the high performance of the CNV-based method on the Peng_et_al_PDAC dataset. The Consensus method was able to retain and emphasize these CNVs as well.

However, the CNV-based method produced weak CNV profiles for the Qian_et_al_Breast samples, which do not support a clear separation between tumor and normal cells, explaining the poor annotation performance with many false positive tumor annotations from normal cell types (as shown in Fig. 3B). On the contrary, while inferring the CNV profiles on the epithelial cells only, the Consensus method was able to detect clear CNV profiles for the same Qian_et_al_Breast samples, supporting tumor vs normal epithelial annotation and avoiding false tumor annotation from other normal cell types. Overall, this shows that the Consensus method is able to produce precise MACE annotations, with clear CNV profiles supporting the malignancy of the annotated tumor cells.

## 4. Discussion

We present a benchmarking study for automated cancer annotation methods using scRNA-seq data. Across 20 cancer datasets covering 9 different cancer types, we compared the state-of-the-art method from the Reference-based category, scATOMIC, and the CNV-based category, SCEVAN. The benchmarking results showed differences in annotation performance depending on the cancer type. Further, we introduced a new Consensus annotation strategy, combining the two previous methods, showing overall superior annotation performance across all cancer types, when compared to applying each method separately.

The Reference-based method can accurately separate epithelial cells from all other cell types. Where in solid tumors, epithelial cells are the main source of malignant cells. However, the Reference-based method cannot further separate normal epithelial cells from those who turned into a malignant state. This limitation is the main cause of the poor annotation performance across cancer types known for strong CNVs like pancreatic cancer.

The CNV-based method can handle cancer types with strong and clear CNVs, producing precise tumor annotations. However, the assumption that all cancer cells undergo copy number aberration is not necessarily valid across all cancer types, where weak CNV profiles are obtained. The final step of the CNV-based method SCEVAN, similar to earlier CNV-based methods, is to cluster cells according to their obtained CNV profiles into two clusters using hierarchical clustering. The cluster with most initial/confident normal cells (providing the baseline normal diploid profile) is consider all normal, while the other cluster is considered tumor. With weak CNV profiles, this final clustering step, forcing the data into two clusters, randomly splits the cells between tumor and normal across all cells in the TME. Moreover, when applied on a completely normal tissue sample, the CNV-based will always produce some tumor annotations, as a result of this forced clustering. However, the Reference-based method will accurately report no tumor cell in the case of a normal tissue sample.

The Consensus annotation method combines the power of each method and overcomes their individual limitations. The Reference-based method ensures accurate annotation of epithelial cells. Also, this ensures to report no tumor cells when applied to a completely normal tissue sample with no epithelial cells. Secondly, applying the CNV-based method only on the epithelial cells compartment avoids wrong tumor annotations across other normal cell types in the TME. Thirdly, it ensures to separate normal epithelial cells, with diploid CNV profiles, from tumor epithelial cells with copy number abnormalities.

As previously mentioned, we focused on the precision score to evaluate the MACE annotation of all methods, to ensure low false positive annotations, i.e. minimal contamination of normal cells mis-annotated as tumor. We argue that a false positive annotation is more harmful than a false negative annotation, with tumor cells being mis-annotated as normal. However, optimizing for high precision may produce a trade-off with lower recall score, indicating higher false negative annotations.

We relied on the authors cell type annotations as the ground-truth for evaluating the MACE annotations. However, it is important to mention that the authors annotations represent a limitation in the current benchmark. As the evaluation presented is dependent and affected by the methodologies used by the authors to generate the original cell type annotations. For example, datasets where authors annotated tumor cells based on CNV-based methods had on average better performance for the CNV-based method in this benchmark.

The Consensus annotation showed precise MACE annotations using scRNA-seq data. Possible improvement can be achieved relying on spatial transcriptomics data, which is growing in the field of single-cell analysis and oncology. With spatial transcriptomics, next to the gene expression data, the spatial distribution of different cells can be used to further inform and improve the final MACE annotation.

Altogether, we present a benchmarking of automated cancer cell annotation methods using scRNA-seq data. We suggest a new Consensus annotation method combining the power of the current state-of-the-art methods and producing precise tumor vs normal cell annotation in the context of solid tumors.

